# IL-33 regulates gene expression in intestinal epithelial cells independently of its nuclear localization

**DOI:** 10.1101/291039

**Authors:** Zhengxiang He, Lili Chen, Glaucia C. Furtado, Sergio A. Lira

**Author notes:** Address correspondence to: Sergio A. Lira, M.D. Ph.D., 1425 Madison Ave, Box 1630 Room 12-20, New York, NY 10029. Z.H. and L.C. contributed equally to this work. **Abbreviations** IBD: inflammatory bowel diseases IEC: intestinal epithelial cells IL-33: Interleukin-33 ILC: innate lymphoid cells LI: large intestine LPL: lamina propria lymphocytes q-PCR: reverse-transcription polymerase chain reaction SI: small intestine Th: T helper cells WT: wild type.

## Abstract

IL-33 is a cytokine found in the extracellular space (mature IL-33) or in the cell nucleus (full-length IL-33). Nuclear accumulation of IL-33 has been reported in intestinal epithelial cells (IEC) during intestinal inflammation and cancer, but a functional role for this nuclear form remains unclear. To study the role of nuclear IL-33 ^in^ IEC, we generated transgenic mice expressing full-length IL-33 ^in^ the intestinal epithelium (*Vfl33* mice). Expression of full-length IL-33 in the epithelium resulted in accumulation of IL-33 protein in the nucleus and secretion of IL-33. Over-expression of full-length IL-33 by IEC did not promote gut inflammation, but induced expression of genes in the IEC and lamina propria lymphocytes (LPL) that correlated negatively with genes expressed in inflammatory bowel diseases (IBD). Because the IL-33 receptor ST2 is expressed by IEC, there was the potential that both the mature and full-length forms could mediate this effect. To specifically interrogate the transcriptional role of nuclear IL-33, ^we^ intercrossed the *Vfl33* mice with ST2-deficient mice. ST2 deficiency completely abrogated the transcriptional effects elicited by IL-33 expression, suggesting that the transcriptional effects of IL-33 on IEC are mediated by its mature, not its nuclear form.

**Highlights:**

- Expression of full-length IL-33 in the epithelium resulted in accumulation of IL-33 protein in the nucleus and secretion of IL-33.
- Full-length IL-33 induced differential gene expression in IEC and LPL that was negatively associated with intestinal inflammatory diseases
- IL-33 regulated gene expression in IEC via its extracellular (mature) form not via its nuclearform.

## 1. Introduction

Interleukin-33 (IL-33), a member of the IL-1 family of cytokines(1), was originally described as a nuclear protein from human high endothelial venules (2). Subsequent studies showed that IL-33 acts as a cytokine, binding a heterodimeric receptor complex consisting of the ST2 receptor (ST2L) and the IL-1R accessory protein. The expression of this heterodimeric receptor has been detected on a variety of inflammatory cells(3, 4), including eosinophils, basophils, macrophages, T helper 2 cells (Th2 cells), regulatory T cells, NK cells, B cells and group 2 innate lymphoid cells (ILC2)(5–7). IL-33 plays a role in the host defense against infection and has been reported to be involved in the pathogenesis of a wide range of diseases (8).

In the gastrointestinal tract, IL-33 is normally expressed by stromal and immune cells, and IL-33 protein has been detected in the nuclei of such cells (9, 10). IL-33 is not normally expressed by epithelial cells, but recent evidence suggests that it can function as a novel epithelial "alarmin"(11), because it can be released as a danger signal by damaged, stressed, or necrotic cells to alert the immune system of a local threat. Epithelial expression of IL-33 has been reported in samples from patients with ulcerative colitis(9, 10,12–15) and cancer(16).

IL-33 is believed to be a dual-function protein, functioning as conventional cytokine via its extracellular form (mature IL-33) or as a transcriptional regulator via its nuclear form (full-length IL-33). Although the molecular mechanism of release and processing of IL-33 are not yet clear(17), it appears that the full-length IL-33 released from injured or necrotic cells is biologically active(11, 18–20), and this bioactivity can be transiently increased several-fold by limited proteolysis of the N-terminal domain (mature IL-33) in inflamed tissue(21, 22) before bioactivity is lost by destruction or oxidization of the C-terminal core tetrahedron structure(23). The N-terminal domain of full-length IL-33 is necessary for nuclear translocation, but it is unclear where it binds to the chromatin and whether it directly regulates gene expression in the intestinal epithelial cells (IEC). In this study, we investigate the biological properties of the full-length IL-33, focusing on its transcriptional properties.

## 2. Materials and methods

### 2.1 Mouse strains

C57BL/6 mice were purchased from The Jackson laboratory (Bar Harbor, ME). ST2^-/-^ mice were generated in our laboratory as described by He et al(16). Mice were maintained under specific pathogen-free conditions. All experiments involving animals were performed following guidelines of the Animal Care and Use Committee of the Icahn School of Medicine at Mount Sinai.

### 2.2 Generation of transgenic mice expressing IL-33 in the intestinal epithelium

The cDNA of IL-33 full-length form was cloned into a pBS-Villin vector that contained a 9kb segment of the mouse villin promoter(24). The pBS-Villin/IL-33 plasmid was verified by sequencing, and the transgene was isolated from the plasmid by restriction enzyme digestion and gel purification. To generate transgenic mice, the transgene was microinjected into C57BL/6 mouse eggs. Identification of the transgenic *Vfl33* mice was done by PCR amplification using the following primers: 5’-ggctgtgatagcacacagga-3’ and 5’-ttcgcctgcggtgctgctgaac -3’.

### 2.3 Enzyme-linked immunosorbent assay

Small pieces of small intestine or colon (5 mm of mid-part) were isolated, rinsed in PBS, weighed, and cultured overnight in 12-well tissue culture plates (Costar) in 1000 μl complete DMEM at 37°C in an atmosphere containing 5% C0_2_. After centrifugation to pellet debris, culture supernatants were transferred to fresh tubes and stored at -80°C. IL-33 was quantified in the supernatant of intestinal explant cultures from *Vfl33* and WT mice by enzyme-linked immunosorbent assay (ELISA) according to standard manufacturer’s recommendations (eBioscience) and the results were normalized to the weight of the intestinal explant.

### 2.4 Reverse-transcription polymerase chain reaction

Total RNA from tissues cells was extracted using the RNeasy mini Kit (Qiagen) according to the manufacturer’s instructions. Complementary DNA (cDNA) was generated with Superscript III (Invitrogen). Quantitative PCR was performed using SYBR Green Dye (Roche) on the 7500 Real Time System (Applied Biosystems) machine. Thermal cycling conditions used were as follows: 50 °C for 2min and 95 °C for 10 min, 40 cycles of 95 °C for 15 s, 60 °C for 1min, followed by dissociation stage. Results were normalized to the housekeeping gene Ubiquitin. Relative expression levels were calculated as 2(^ct(Ubiquitin)-ct(25)^), Primers were designed using Primer3 Plus software(26).

### 2.5 Histology and immunofluorescence staining

Tissues were dissected, fixed in 10% phosphate-buffered formalin, and then processed for paraffin sections. Five-micrometer sections were stained with hematoxylin and eosin (H&E) for histological analyses. For immunofluorescence staining, five-micrometer sections were dewaxed by immersion in xylene (twice for 5 minutes each time) and hydrated by serial immersion in 100%, 90%, 80%, and 70% ethanol and PBS. Antigen retrieval was performed by microwaving sections for 20 minutes in Target Retrieval Solution (DAKO). Sections were washed with PBS (twice for 10 minutes each time), and blocking buffer (10% BSA in TBS) was added for 1 hour. Sections were incubated with primary antibody in blocking buffer overnight at 4°C. After washing, conjugated secondary Abs were added and then incubated for 35 min. Cell nuclei were stained using 4’,6-Diamidino-2-Phenylindole (DAPI). The slides were next washed and mounted with Fluoromount-G (Southern Biotech). Images were captured using a Nikon fluorescence microscope. Colocalization was performed with ImageJ and the colocalization finder plug-in.

### 2.6 Western blot analysis

Intestine were opened longitudinally and thoroughly washed in PBS and then homogenized in ice-cold lysis buffer (100 mM Tris-HCl, pH 6.8, 4% SDS, 20% glycerol, 200 mM (β-mercaptoethanol, 1 mM phenylmethylsulfonyl fluoride and 1 μg/mL aprotinin). Lysates were then centrifuged at 12 000g for 15 min to remove insoluble cell debris. Protein content was quantified using the Bio-Rad protein assay (Bio-Rad) and 15 μg of protein was separated by SDS-PAGE and transferred onto polyvinylidene difluoride membranes. The membrane was blocked for 1 h in buffer (TBS, 5% milk, 0.1% Tween 20) and then incubated with the primary antibody (Rat Anti-Mouse IL□33 Monoclonal Antibody) (Catalog # MAB3626, R&D Systems) in dilution buffer (TBS, 5% bovine serum albumin, 0.1% Tween 20) overnight at 4 °C. The membrane was then washed three times with wash buffer (TBS, 0.1% Tween 20), incubated with Rat IgG HRP-conjugated Antibody (Catalog # HAF005, R&D Systems) and visualized with the enhanced chemiluminescent detection system (Amersham Biosciences).

### 2.7 Isolation of IEC and LPL

Intestines were opened longitudinally and thoroughly washed in PBS. The intestine was then incubated in 30 ml PBS containing 1 mM dithiothreitol (DTT) on room temperature for 15 min. The intestine was then removed and briefly washed in PBS and incubated in 25 ml PBS containing 5.2 mM ethylenediaminetetraacetic acid (EDTA) at 4 °C at 200 RPM for 30 min. The cells were then subjected to 30 sec vigorous shaking and the tissue removed. The cells were then centrifuged at 1000 g for 5 min at 4 °C, washed in PBS containing 10% FBS and spun for a further 5 min at 4 °C at 1000 g. These cells constituted the IEC population. To isolate lamina propria lymphocytes (LPL), the remaining tissues were performed as described before(16).

### 2.8 Cell sorting

Cell pellets were first pre-incubated with anti-mouse CD16/CD32 for blockade of Fc γ receptors, then were washed and incubated for 30 min with fluorescent conjugated antibodies against CD45 and EpCAM in a total volume of 500 µl PBS containing 2 mM EDTA and 2*%* (vol/vol) fetal bovine serum. DAPI (Invitrogen) was used to distinguish live cells from dead cells during cell sorting. Stained lECs (DAPI^-^CD45^-^Epcam^+^) and LPL (DAPI^-^CD45^+^) were purified with a MoFlo Astrios cell sorter (DakoCytomation). Cells were > 95% pure after sorting.

### 2.9 Microarray analysis

Total RNA from the sorted intestinal CD45^+^ cells from WT and *Vfl33* mice was extract using RNeasy Micro Kit (Qiagen). Microarrays were done and analyzed as described before(27, 28). In order to analyses the pathways that the differentially expressed genes are involved in, KEGG pathway enrichment analyses were performed using ClueGo(29, 30). A cut-off of 0.4 was set for kappa score and terms including at least 3 genes were retrieved.

### 2.10 RNA-seq

Following cell sorting into Trizol LS reagent, samples were shipped on dry ice to the Center for Functional Genomics and the Microarray & HT Sequencing Core Facility at the University at Albany (Rensselaer). Total RNA from sorted cells (3 -9 X10^5^cells) was extracted using the RNeasy micro Kit (Qiagen) with an on-column DNAse digestion step included according to the manufacturer’s instructions. RNA quality was assessed using the Nanodrop (Thermo Scientific) and Bioanalyzer Total RNA Pico assay (Agilent). Total RNA with a RNA integrity number (RIN) value of 7 or greater was deemed of good quality to perform the subsequent protocols. 100 pg of total RNA was oligo-dT primed using the SMART-Seq v4 Ultra Low Input RNA Kit (Clontech) and resulting the cDNA was amplified using 15 cycles of PCR. The double stranded cDNA (dscDNA) was purified using AMPure XP magnetic beads and assessed for quality using the Qubit dsDNA HS assay and an Agilent Bioanalyzer high sensitivity dscDNA chip (expected size ~600bp-9000bp). The Illumina Nextera XT kit was used for library preparation wherein 125 pg dscDNA was fragmented and adaptor sequences added to the ends of fragments following which 12 cycles of PCR amplification was performed. The DNA library was purified using AMPure XP magnetic beads and final library assessed using Qubit dsDNA HS assay for concentration and an Agilent Bioanalyzer high sensitivity DNA assay for size (expected range ~600-740bp). Library quantitation was also done using a NEBNext Library Quant kit for Illumina. Each library was then diluted to 4nM, pooled and denatured as per standard Illumina protocols to generate a denatured 20 pM pool. A single end 75bp sequencing was performed on the Illumina Nextseq 500 by loading 1.8 pM library with 5% PhiX on to a 75 cycle high output flow cell. The RNAseq data was checked for quality using the Illumina FastQC algorithm on Basespace.

### 2.11 Transcriptome analysis

RNA-Seq data from IECs was mapped to the mouse reference genome (UCSC/mm10) using Tophat version 2.1.0(31). Gene-level sequence counts were extracted for all annotated protein-coding genes using htseq-count version 0.6.1 (32) by taking the strict intersection between reads and the transcript models associated with each gene. Raw count data were filtered to remove low expressed genes with less than five counts in any sample. Differentially expressed genes between groups were analyzed using Bioconductor EdgeR package version 3.10.2 Bioconductor/R(33, 34). Statistically significant differentially expressed genes between groups (Q < 0.05) were selected in gene-wise log-likelihood ratio tests that were corrected for multiple testing by Benjamini and Hochberg FDR.

### 2.12 NextBio

Meta-analysis was conducted by NextBio (www.nextbio.com) (35). The gene list (*Vfl33* vs WT) from LPL Microarray results and IEC RNA-seq results were used as input to query a collection of individual biosets in NextBio database. The NextBio application "disease atlas" was used. The association score based on statistical significance across different diseases (100 to the most significant inside the diseases).

### 2.13 Statistics

Differences between groups were analyzed with nonparametric Mann-Whitney test. For the comparison of more than two groups a one-way ANOVA followed by a Bonferroni multiple comparison tests were performed. All statistical analyses were performed using GraphPad Prism 5 software.

## 3. Results

### 3.1 Generation of transgenic mice expressing nuclear IL-33 in the epithelium

To investigate the function of full-length IL-33 in epithelial cells, we cloned the cDNA encoding the full-length form of IL-33 downstream of the villin promoter (Fig. 1A). The transgene was injected into mouse eggs and 3 transgenic lines were derived. These animals are referred to as *Vfl33* mice (Fig. 1A). The *Vfl33* mice were healthy and reproduced normally. To examine IL-33 expression and select a line for further studies, we extracted RNA from the small and large intestine of control and transgenic mice. As expected, we detected increased expression of IL-33 mRNA in the small intestine and large intestine in the *Vfl33* transgenic mice compared with their littermate control WT mice (Fig. 1B). To examine expression of IL-33 protein, we performed ELISA in the gut extracts and found that IL-33 production in the supernatant was elevated in the intestine of transgenic mice compared to WT mice (Fig. 1C). Finally, we examined the cellular expression of IL-33. Because we expressed the full-length form of IL-33, we expected that it should be located in the nucleus rather than the cytoplasm. Immunostaining of intestinal sections showed that IL-33 immunoreactivity was indeed detected in the nucleus of transgenic, but not control intestinal epithelial cells (Fig. 1D). Similar qualitative results were obtained by analysis of animals in the 3 transgenic lines and line 2 was selected for further analysis.

**Fig. 1.**
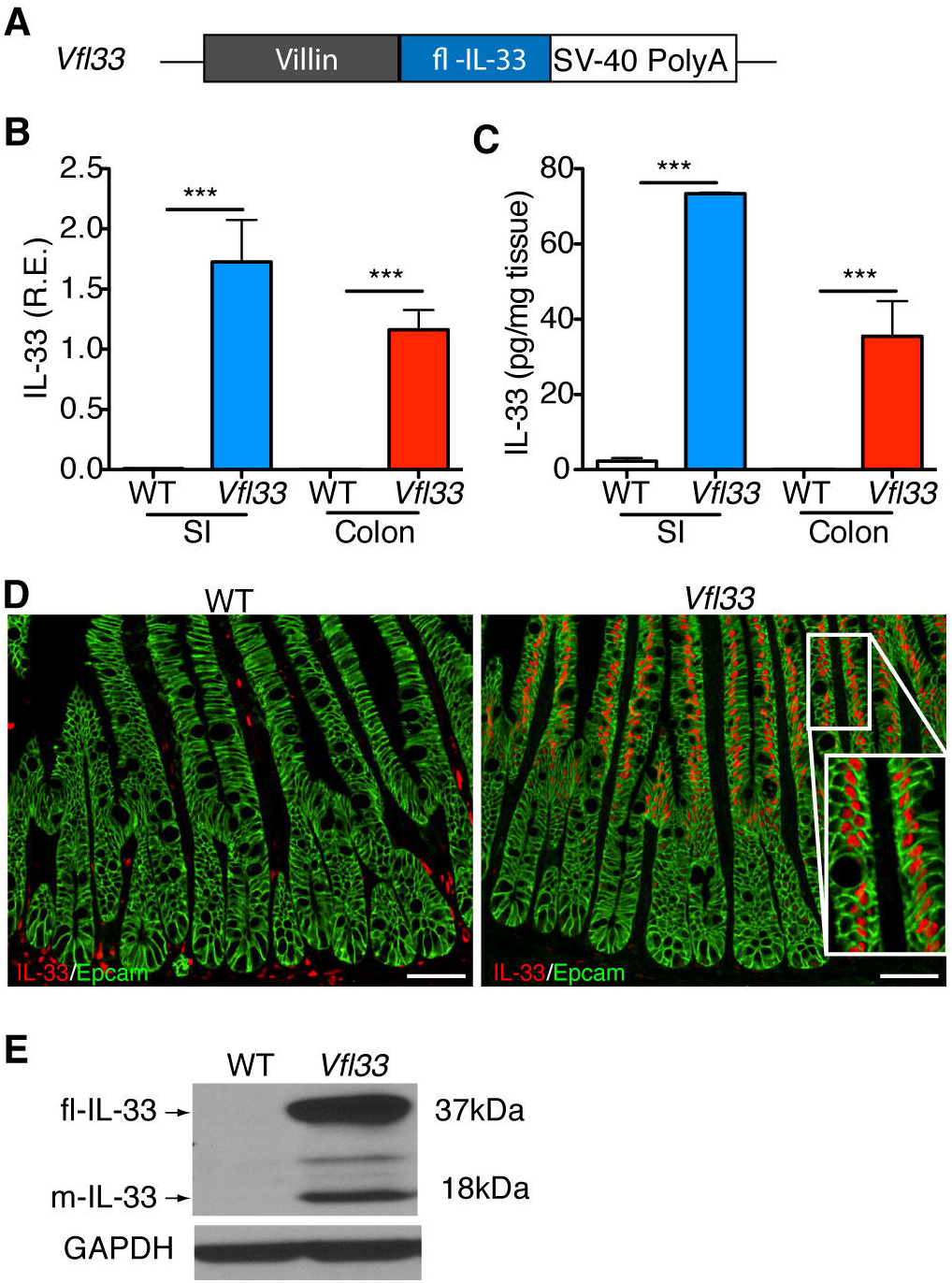
Generation of transgenic mice expressing full-length IL-33 in the gut epithelium. (A) Scheme for generation of *Vfl33* mice. A transgene encoding the IL-33 full-length form under the control of the murine villin promoter (9kb) was used to generate *Vfl33* mice. (B) Relative expression levels of IL-33 mRNA were analyzed by qPCR in the small intestine (SI) and colon of wild-type (WT) and *Vfl33* mice. Data were normalized to the expression levels of the Ubiquitin transcript. Means ± s.e.m., n = 6 per group. ***P < o.oo1, one-way ANOVA. (C) Enzyme linked immunosorbent assay of IL-33 in the gut explants from WT and *Vfl33* mice. Data were normalized to the weight of the intestine explant. Means ± s.e.m., n = 4 per group. ***P < 0.001, one-way ANOVA. (D) Immunofluorescence staining for IL-33 (red) in the gut of WT and *Vfl33* mice. Notice that transgenic expression of IL-33 in the nucleus of intestinal epithelial cells (IEC) in *Vfl33* mice. IEC stained with an anti-Epcam antibody (green). Scale bars, 5o μm. Inset shows higher magnification of the boxed area. (E) Western blot analysis of IL-33 expression in the gut of WT and *Vfl33* mice. The GAPDH protein levels were used as protein loading controls. The expected molecular sizes for the full-length and mature forms of IL-33 are 37 and 18 kDa, respectively.

Although the molecular mechanism of release and processing of IL-33 are not yet clear(17), the studies done so far show that the full-length IL-33 released from injured or necrotic cells is biologically active(11, 18–20), and that its bioactivity can be transiently increased several-fold by limited proteolysis of the N-terminal domain (mature-IL-33) in the tissue(21, 22). We next examined whether mature IL-33 could be detected in the intestine of *Vfl33* mice. To do so, we performed Western blot to analyze the different molecular species of IL-33 in the intestine. We found that the predominant form of IL-33 in the extracts was the full-length form (37kD band), but that a 18kD (mature IL-33) was also present (Fig. 1E). Together the results indicate that the transgenic IL-33 was correctly targeted to IEC of both small and large intestine, and that it accumulated in the nucleus. In addition they suggest that the processed IL-33 mature form was also produced, although at lower concentration.

### 3.2 Over-expression of full-length IL-33 by IEC does not promote intestinal inflammation

To examine the potential impact of epithelial-specific IL-33 on the intestinal inflammation, we performed histological analysis of the gut in *Vfl33* mice. We found that there was no inflammation in the gut of transgenic mice (Fig. 2A). Consistent with this, FACS analysis of LPL showed that there were no differences in the total number of CD45^+^ cells between *Vfl33* and WT mice (Fig. 2B). The results suggested that over-expression of full-length IL-33 by IEC did not promote gut inflammation.

**Fig. 2.**
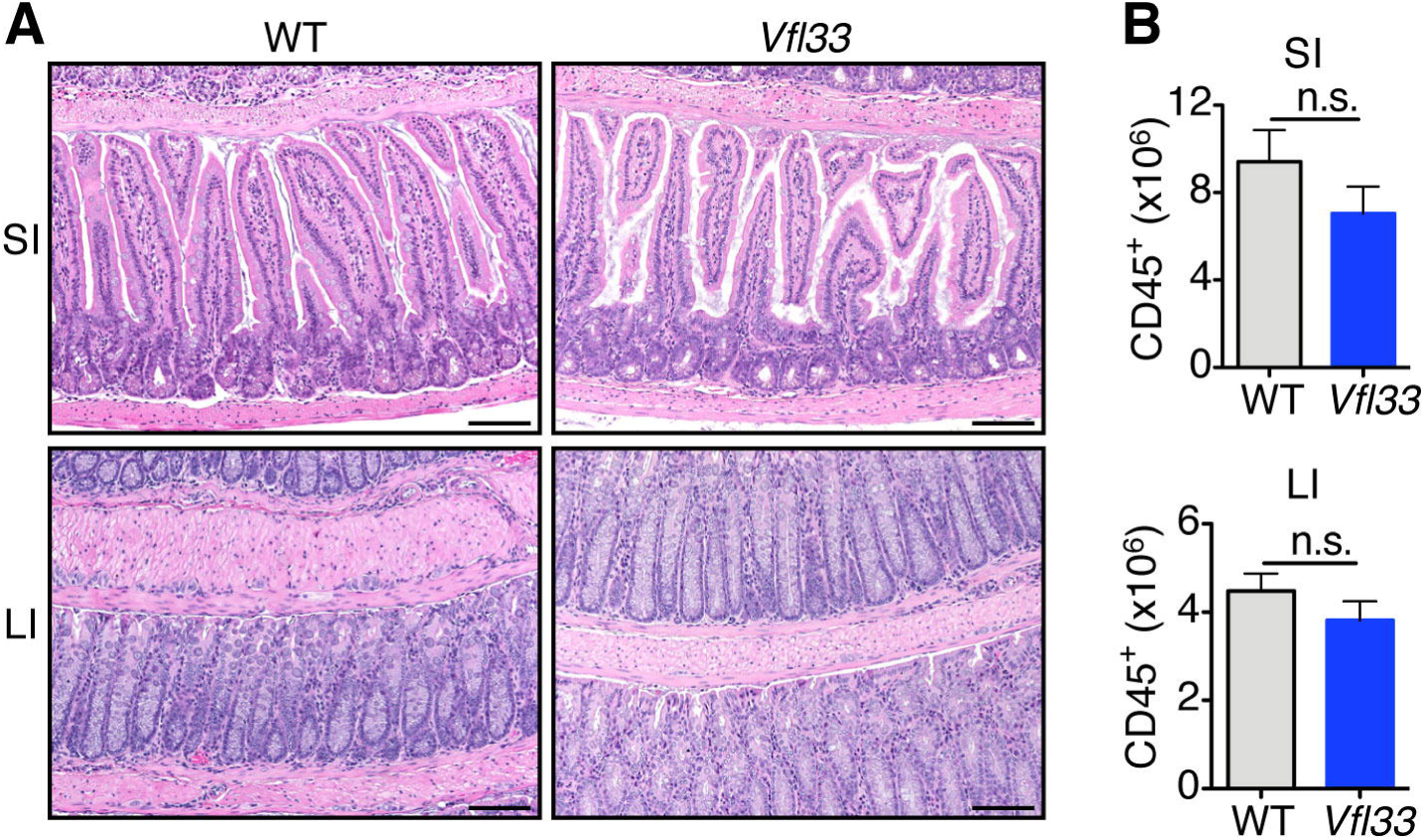
Over-expression of full-length IL-33 by IEC does not promote gut inflammation. (A) Representative H&E-stained intestinal sections of *Vfl33* mice. Scale bars, 50 μm. (B) Total CD45^+^ cells in the gut of *Vfl33* and WT mice (n=6 mice/group). N.s, non significant; by Mann-Whitney test.

### 3.3 Epithelial-derived IL-33 triggers gene expression in lamina propria leukocytes

To examine if IL-33 expression affected intestinal immune cells, we sorted lamina propria CD45^+^ cells from both groups to perform microarray analysis. The results show that 105 genes were upregulated and 58 genes were downregulated in *Vfl33* mice compared to WT mice (Fig. 3A &3B). It has been reported that IL-33 can enhance the primary differentiation of CD4^+^ Th1, Th2, and Treg (36). We found that expression of full-length IL-33 was associated with increased Th2 immune response in the gut, consistent with a signature of up-regulated Th2 transcription factor (*Gata3)* and Th2 cytokines (*Il4* and *Il13*) by quantitative PCR (q-PCR) (Fig. 3C). In addition, expression of *retnla* and *retnlb,* which are induced in Th2 environment(37), was also up-regulated in the gut of *Vfl33* mice (Fig. 3C).

**Fig. 3.**
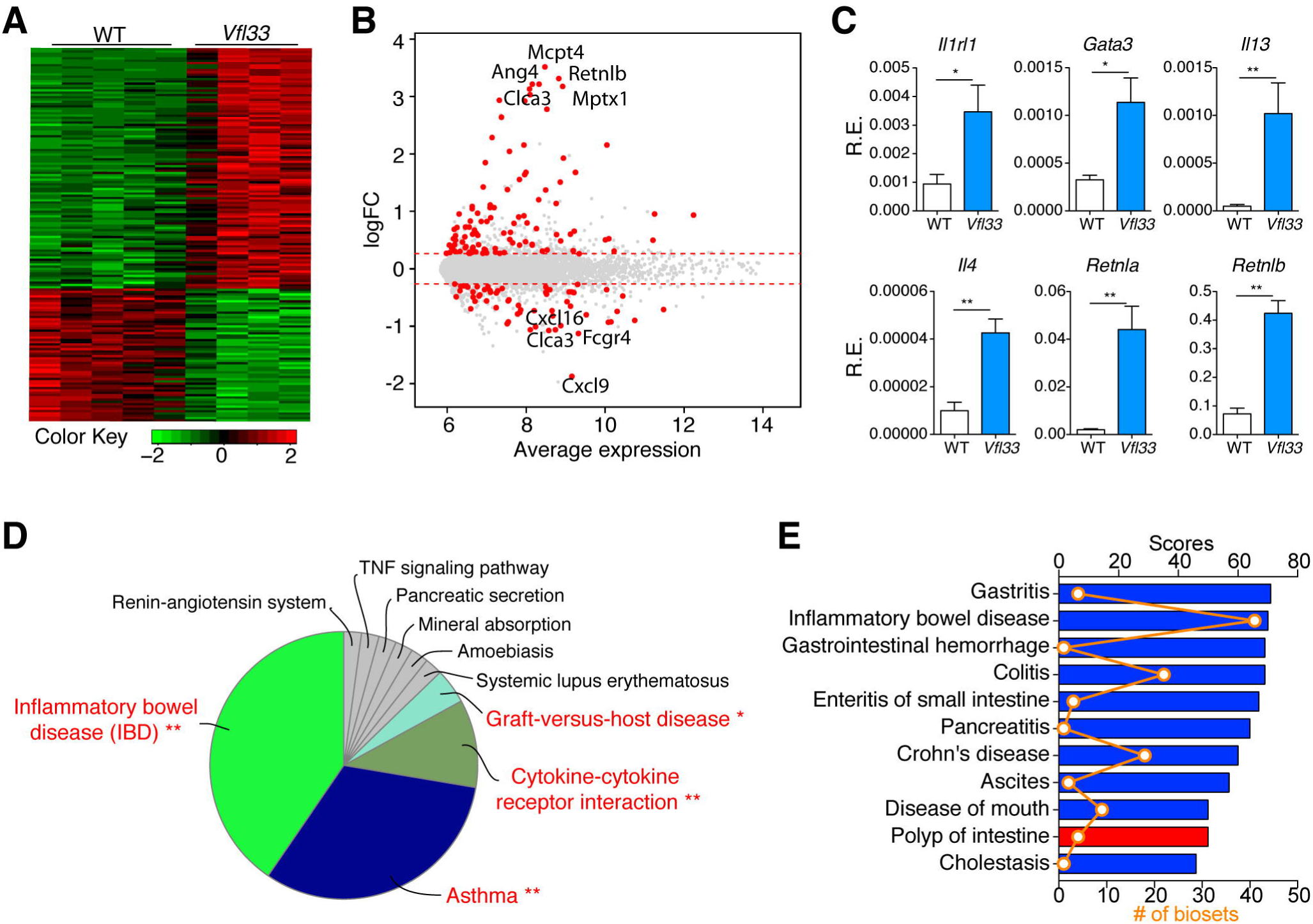
Epithelial-derived IL-33 triggers gene expression in lamina propria leukocytes. (A-B) Leukocytes (CD45^+^) isolated from *Vfl33* and WT LPL were analyzed by microarray (n= 4-5/group). (A) Quantile-normalized expression values were analyzed using a paired design and filtered for Q < 0.05 and -1.2 > fold change > 1.2. Z score-normalized data were subjected to hierarchical clustering. (B) Plot of logFC (log fold change) versus mean expression of all detected transcripts (gray) and significant genes (red). (C) Quantitative PCR analysis of the selected Th2 immune response associated genes in the gut of *Vfl33* and WT mice (n= 5-7/group). Data were normalized to the expression levels of the Ubiquitin transcript. Means ± s.e.m., *P < 0.05, **P < o.o1, Mann-Whitney test. (D) The pathway enriched among total 163 differential expressed genes in the *Vfl33* versus WT LPL by KEGG analysis. (E) The mRNA expression profile of *Vfl33* LPL vs WT LPL correlated with various digestive system diseases by using NextBio meta-analysis program. The correlated diseases are shown with the corresponding significant scores as well as the number of biosets within each disease. Columns are colored according to correlation with query: positive correlation (red) and negative correlation (blue).

Pathway analysis of differentially expressed mRNAs is designed to provide insight into the cell pathways associated with these genes. Pathway analysis of 163 differential expressed genes in CD45^+^ cells of *Vfl33* mice showed that "inflammatory bowel disease" was the top pathway (Fig. 3D), suggesting that epithelium expressed IL-33 could have a role in the IBD. In addition, we used the gene list (*Vfl33* vs WT) from the LPL microarray as input to query a collection of individual biosets in NextBio database. The NextBio application "disease atlas", which focuses on digestive system disease, was used for analysis. We found that differentially expressed *Vfl33 genes* negatively correlated with genes expressed in inflammatory bowel disease, Crohn’s disease, enteritis of small intestine and colitis (Fig. 3E). Together, the pathway analyses and the NextBio meta-analysis suggest that epithelium-derived IL-33 regulates gene expression, and that many of these differentially expressed genes negatively correlate with genes expressed in IBD.

### 3.4 Full-length IL-33 regulates gene expression in the epithelium

To investigate the function of IL-33 in the epithelium, we sorted intestinal epithelial cells from *Vfl33* mice and WT mice, extracted mRNA and preformed RNA sequencing. The results showed that 103 genes were upregulated and 52 genes were downregulated in transgenic IEC compared to controls (Fig. 4A). Since the IL-33 regulated genes in the lamina propria leukocytes were associated with IBD, we asked next if the differentially expressed genes of the intestinal epithelial cells also correlated with IBD. To do so we filtered the differentially expressed genes in the IEC RNA-seq results using the disease category locator of the NextBio software. The results indicated that the differentially expressed genes shared high similarity with genes associated with digestive system disease (Fig. 4B). Notably, differentially expressed genes were negatively correlated with intestinal inflammatory diseases such as enteritis of small intestine, IBD, colitis and Crohn’s disease (Fig. 4B). Taken together, the transcriptomic analyses of both epithelial cells and leukocytes suggest a protective role of epithelial-derived IL-33 in intestinal inflammation.

**Fig. 4.**
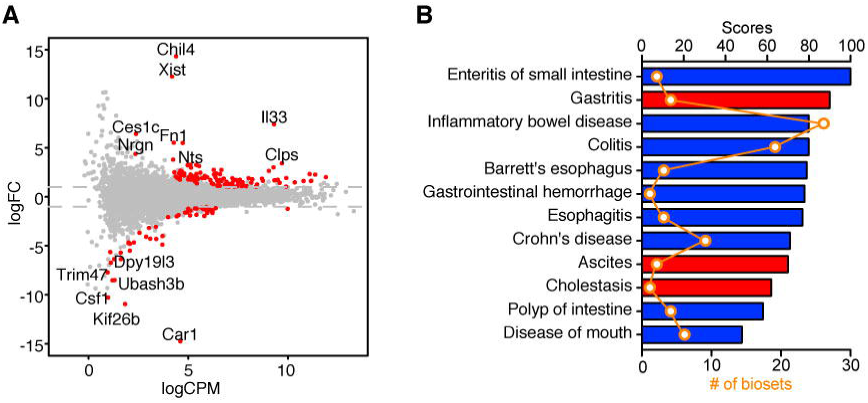
Differential expression of epithelial genes triggered by IL-33 correlates with a subset of genes involved in diseases of the digestive system. (A) *Vfl33* and WT IEC were analyzed by RNA-Seq/edgeR using a paired design (n = 3-4/group). Plot of logFC (log fold change) versus logCPM (log counts per million) of all detected transcripts. Points are colored according to expression status: non-significant genes (gray) and significant genes (155 genes; Q < 0.05 and -2 > fold change > 2; red). (B) NextBio meta-analysis showed that the mRNA expression profile of *Vfl33* IEC vs WT IEC negatively correlated with various digestive system diseases. The correlated diseases are shown with the corresponding significant scores as well as the number of biosets within each disease. Columns are colored according to correlation with query: positive correlation (red) and negative correlation (blue).

### 3.5 IL-33-induced gene regulation in IEC is not dependent on its nuclear localization

IL-33 is hypothesized to be a dual-function protein, functioning as a conventional cytokine and/or as a transcriptional regulator. To investigate if the nuclear form of IL-33 directly regulated transcription, we expressed the IL-33 full-length form and simultaneously deleted expression of its receptor ST2. To do so, we crossed IL-33 transgenic mice (*Vfl33*) with ST2 deficient mice *(ST2^-/-^)(16)* to generate *Vfl33 ST2^-/-^*mice. First, we confirmed that shutdown the IL-33/ST2 signaling did not change the nuclear location of full-length IL33 in *Vfl33 ST2^-/-^* mice (Fig. 5A). We then sorted IEC from WT, *Vfl33, ST2^-/-^* and *Vfl33 ST2^-/-^* mice, extracted RNA and preformed RNA-seq. To our surprise, WT, *ST2*^*-/-*^ and *Vfl33* S*T2*^*-/-*^ clustered together, and apart from *Vfl33* by principal component analysis (Fig. 5B). Further analysis shown that with exception of IL-33, there were no differences in gene expression between IEC from *Vfl33 ST2*^*-/-*^ and those from WT mice (Fig. 5C). In addition, there were no differences in gene expression between IEC of WT and *ST2*^*-/-*^ mice (Fig. 5B). As shown in Fig. 5D, none of the differentially expressed genes between *Vfl33* and WT mice were differentially expressed in *Vfl33 ST2*^*-/-*^ mice. Thus, the RNA-seq analyses do not support a role of nuclear IL-33 in the regulation of gene expression in the IEC. All together, the results suggested that IL-33 regulates gene expression in the epithelium via its interaction with epithelial-expressed ST2.

**Fig. 5.**
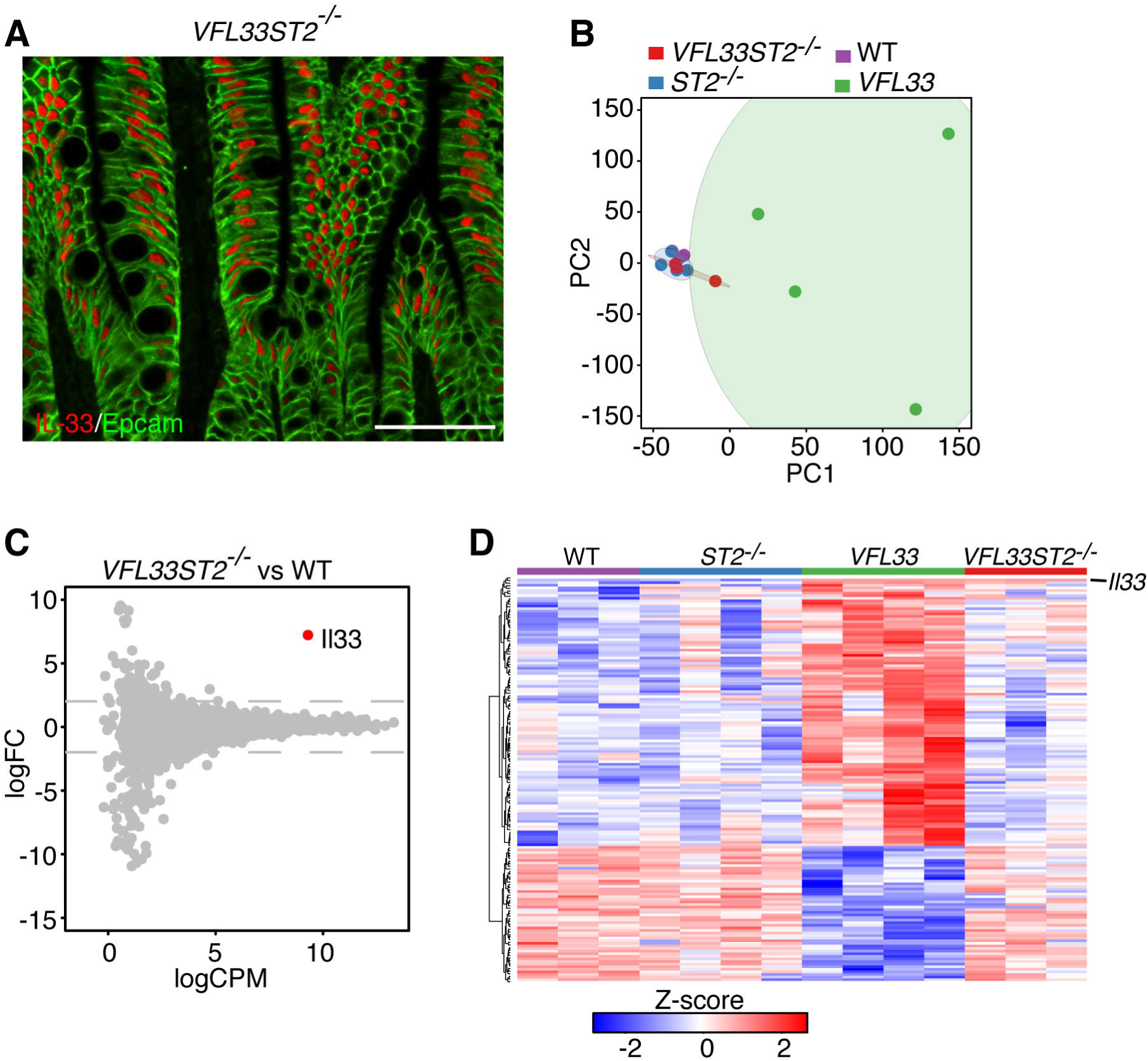
Genetic deletion of ST2 in Vfl33 mice abrogates IL-33 induced gene expression in IEC. (A) Immunofluorescence staining for IL-33 (red) in the intestine of *Vfl33 ST2*^*-/-*^ mice. Scale bars, 50 μm. (B) Principal component analysis of RNA-seq expression data from all biological replicates of IEC from WT, *Vfl33*, ST2^-/-^and *Vfl33 ST2*^-/-^ (n = 3-4/group). (C) *Vfl33 ST2*^*-/-*^and WT IEC were analyzed by RNA-Seq/edgeR using a paired design (n = 3-4/group). Plot of logFC (log fold change) versus logCPM (log counts per million) of all detected transcripts. Points are colored according to expression status: non-significant genes (gray) and significant gene (*II33* gene; Q < 0.05; red). (D) Heat map depiction of all differentially expressed genes between WT and Vfl33 in all groups (WT, *Vfl33, ST2*^-/-^ and *Vfl33 ST2^-/-^).* Red indicates increased and green indicates decreased expression in *Vfl33* IEC compared with WT IEC.

## 4. Discussion

IL-33 has been detected in the nucleus of epithelial cells in intestinal inflammation and cancer, but whether this nuclear form of IL-33 contributes to transcriptional regulation, is a matter of debate. In this study, we generated transgenic mice to investigate the role of IL-33 on IEC (Fig. 1). We found that the bulk of the IL-33 protein present in intestinal extracts of these animals corresponded to the full-length form, but also detected the mature, processed form. Immunostaining experiments documented the nuclear accumulation of IL-33 in the IEC. Comparison of gene expression profiles of sorted IEC from WT and *Vfl33* mice showed a marked difference, clearly supporting a role for IL-33 in regulating IEC gene expression (Fig. 3A). The precise effect of epithelial-derived IL-33 to the gut inflammatory conditions has remained unclear. Experimental data obtained using different animal models of intestinal inflammation have produced conflicting results(38, 39), with IL-33 having both pro-inflammatory and anti-inflammatory effects. Previous studies done by our group have shown that expression of mature IL-33 by IEC does not promote gut inflammation(16), suggesting that epithelium derived mature IL-33 does not display pro-inflammatory properties. In line with this observation, over-expression of full-length IL-33 by IEC reported here also does not promote gut inflammation (Fig. 2). Meta-analysis of genes differentially expressed in *Vfl33* mice in the IEC negatively correlated with intestinal inflammatory diseases such as enteritis of small intestine, IBD, colitis and Crohn’s disease (Fig. 4), suggesting a protective role of epithelial-derived IL-33 in the intestinal inflammation.

We suggest that the expression of IL-33 by IEC, contributes a protective effect on the acute inflammation that ensues from damage of the gut epithelium. Therefore, IL-33 production may serve as a counterregulatory response to inflammation. The protective effects could derive from a direct cytokine role of IL-33 on IEC, leading to expression of genes that are negatively associated with inflammation as described above. This protective ability may not be only related to the effects of IL-33 signaling on IEC. Release of the mature IL-33 could also affect immune cells. Indeed, our microarray analyses indicate that epithelium-derived IL-33 differentially regulated expression of several genes in lamina propria leukocytes. One of the most upregulated genes in lamina propria leukocytes was the IL-33 receptor ST2. Previous work from our lab has shown that epithelial-derived IL-33 induces expansion of ST2^+^ Treg cells in the intestine(16). Tregs induced by IL-33 signaling could ameliorate inflammation(40). Other genes to prominently expressed by the *Vfl33* leukocytes were those involved in the Th2 response in the gut (Fig. 3). Indeed, induction of Th2 responses may be of benefit when mucosal inflammation is mediated through Th1 or Th17 pathways(41). These results add to the clinical evidence that administration of helminths as a therapeutic option in IBD, as helminths may exert their anti-inflammatory effect through the induction of specific Th2 cytokines(42, 43). Epithelial-derived IL-33 could directly act on Th17 cells to help them to acquire immunosuppressive phenotype(44). In addition, IL-33 induced M2 macrophages has been reported in the contribution of attenuation of colitis(45, 46). Th2 cytokines also appear to influence resolution of inflammation by inducing polarization of macrophages to M2 macrophages (45). Finally, the genes regulated in lamina propria leukocytes by IL-33 negatively correlated with genes associated with intestinal inflammatory diseases (Fig. 3).

Most of the studies on the function of the nuclear function of IL-33 have been performed in endothelial cells(47). It has been shown that nuclear IL-33 can bind to the acidic pocket formed by histones H2A and H2B (48, 49), and to bind to the transcriptional repressor histone methyltransferase (50), but it is unclear if this physical association has functional properties. Some studies have reported that IL-33 can affect nuclear factor-kB (NF-kB) activity in a gene dependent manner. Binding of IL-33 to the NF-kB p65 subunit in the nucleus reduces p65-triggered gene expression to dampen the production of proinflammatory cytokines(51). However, others have reported that nuclear IL-33 could bind to the promoter region of p65, positively regulating its transcriptional activity in endothelial cells(52). Thus, while there is evidence for binding of IL-33 to nuclear proteins in endothelial cells, its ability to regulate gene transcription remains controversial.

Here we provide direct evidence that IL-33 can regulate gene expression in the epithelium. Under these conditions IL-33 could affect gene regulation by acting directly in the nucleus, or by autocrine regulation via ST2, which is expressed by epithelial cells(14), or both. To discriminate among these possibilities, we forced IL-33 expression in epithelial cells and ablated expression of ST2 in all cells, including epithelial cells. Surprisingly, the only gene to be upregulated in these IEC in these conditions was the transgenic full-length, nuclear form of IL-33. The results indicate that IL-33 induced transcriptional changes via autocrine activation of ST2 and suggest that there are no transcriptional activities associated with its nuclear form in IEC (Fig. 5). We cannot formally rule out that the nuclear form of IL-33 controls microRNA transcription, because our methods for RNA preparation did not enrich for microRNAs. The impact of any increased miRNA expression, however, would be in the levels of mRNA transcripts, but no changes were detected when the transcriptomes of WT and *Vfl33 ST2*^*-/-*^ IEC were compared. We would thus posit that there are no transcriptional activities associated with the nuclear form of IL-33 in IEC. Our transcriptional studies in IEC corroborate proteomic studies done by Girard’s group using endothelial cells(53). Using RNA silencing strategies they showed that in endothelial cells IL-33 acts as a cytokine, not as a nuclear factor, regulating gene expression (53). The actual function of the nuclear form may be related to control the availability of the mature form in circulation. Elegant studies done by Bessa et al. (54) show that in the absence of nuclear localization, IL-33 is released into the circulation leading to widespread non-resolving inflammation that culminates in the death of the animal(54). Therefore, the main purpose of IL-33 nuclear localization and chromatin association may be the regulation of its potent extracellular (mature IL-33) cytokine activity(53), not control of transcription, as demonstrated here.

## 5. Conclusion

In summary, expression of full-length IL-33 in the epithelium resulted in accumulation of IL-33 protein in the nucleus and secretion of IL-33. Accordingly, expression of full-length IL-33 in the epithelium promoted expression of genes in the neighboring lamina propria leukocytes and in epithelial cells. The gene program activated by IL-33 in these cells suggests that this molecule has a role in resolution of the inflammatory response. The transcriptional program elicited by expression of full length IL-33 was promoted by its mature, processed, form via binding to its receptor ST2, not by its nuclear form in intestinal epithelial cells.

## Author contributions

L.C. and Z.H. designed study, did experiments, analyzed data and wrote the manuscript; G.C.F and S.A.L designed study, analyzed data and wrote the manuscript. All authors reviewed the manuscript.

## Conflict of interest

The authors declare no competing financial interests.

## Grant Support

This work was supported in part by the SUCCESS (Sinai Ulcerative Colitis Clinical, Experimental and System Studies) grant from the Bacchetta Foundation. Funds for the generation of the *Vfl33* mice were provided by Janssen Pharmaceuticals. Lili Chen received a Research Fellowship Award (327362) from the Crohn’s & Colitis Foundation of America (CCFA).

## Acknowledgments

We thank all members of the Lira Lab for their support. We thank Claudia Canasto-Chibuque for colony maintenance, Chao Yang for Western Blot and Dr. Kevin Kelley and the Mouse Genetics Shared Research Facility for assistance in generation of transgenic mice.

